# Open Access publishing in Medicine: lights and shadows

**DOI:** 10.1101/2021.11.08.467687

**Authors:** Nicola Bernabò, Luca Valbonetti, Alessandra Ordinelli, Rosa Ciccarelli, Barbara Barboni

**Affiliations:** Faculty of Bioscience and Agro-Food and Environmental Technology, University of Teramo, Teramo, Italy; National Research Council – Institute of Biochemistry and Cell Biology, Rome, Italy

**Keywords:** Open Access, Medicine, bibliometrics, Scopus, citation advantage

## Abstract

The widespread use of internet has had enormous consequences in changing the way of accessing to scientific literature in all domains of knowledge and, in particular, in medicine. One of the most important related factors is the idea of making research output freely available: the so-called Open Access (OA) option. OA is considered very important in spreading of knowledge, breaking down barriers in benefit of research, and increasing the impact of research outputs within the scientific community.

Here, we carried out a comparison between Non-Open Access (NOA) and OA medical Journals in terms of growing rate, geographical distribution, and the impact on scientific community.

We collected the bibliometric data on the scientific Journals indexed in Scopus starting from 2001 to 2016 published either as NOA or OA. Then, we analysed the number of Journals, their geolocalization, their impact on the scientific community, and the parameters as SJR, *H* index, and cites for document (2 years).

As a result, we found that while the number of NOA Journals is virtually stable, that of OA is dramatically increasing, with a growing rate higher than 400% in 2016. Then, the majority of OA Journals are published in developing Countries (Brazil, India, South Africa, South Korea, New Zealand, Serbia, and Poland) and their impact within researchers is lower compared to the NOA Journals.

In conclusion, our data provide an updated and unprecedented picture of OA adoption in medical field, with its lights and shadows.

**Article Highlights:**

- We are seeing in medicine a huge increase in the number of Journals that use OA;
- This trend is mainly due to the increase in publishing Journals in Countries with emergent economies;
- The use of OA for Journals in the medical field, did not guarantee a vantage in term of bibliometric parameters.

**MCS:** 92C99

**JEL:** I30

## Introduction

Widespread public access to the World Wide Web, possible nowadays with negligible costs for a large number of people (at least in developed Countries), has deeply changed the way of accessing to scientific knowledge, with enormous consequences in the spread of information among scientists and between scientists and the lay public. In this context, starting from the late 90’s, the issue free access to the scientific literature has been raised, the so-called open access (OA), possible available so far only to scientific Journal subscribers.

More precisely, Open access (OA) is defined the access, to research outputs free from all access and many use restrictions (https://opensource.com/resources/what-open-access). It is possible to distinguish different forms of OA. *Gratis* OA: the online access free of charge, and *libre* OA, the online access free of charge plus various additional usage rights, usually under the Creative Commons licenses. In addition, there are possible Golden OA or Green OA options. Golden OA is provided by Journals that make the research output immediately available on line and generally requires the payment of a publication fee (from hundreds to thousands of dollar), green OA has no cost and is provided by researchers who self-archive their own research outputs in a public repository (such as institutional and central repositories). There are several important reasons related to the adoption of OA. Firstly, it allows the spreading and sharing of knowledge in developing countries, where research Institutions often find it difficult to pay for subscriptions required to have access to scientific literature. In other words, ideally, all researchers (but also journalists, politicians, civil servants, or interested laypeople) could benefit from OA, being able to dispose of scientific knowledge without barriers or restrictions. Then, several funding Agencies, either at international or national level, explicitly and mandatorily require that researchers publish their research output in OA Journals. For instance, EU in the H2020 program requires that all research outputs be published with OA, with the aim of “building on previous research results (improved quality of results), encouraging collaboration and avoiding the duplication of effort (greater efficiency), speeding up innovation (faster progress to market means faster growth) and involving citizens and society (improved transparency of the scientific process)”.^1^. The same policy has been adopted by the Italian Ministry of Educations, University and Research (MIUR) for the research outputs from projects funded by the Projects of Relevant National Interest (PRIN) programme (http://attiministeriali.miur.it/anno-2016/novembre/dd-07112016-(1).aspx). In addition, important Professional Organizations encourage the use of OA such as the International Mathematical Union that in 2001 stated: “Open access to the mathematical literature is an important goal” and encouraged them to “[make] available electronically as much of our own work as feasible” to “[enlarge] the reservoir of freely available primary mathematical material, particularly helping scientists working without adequate library access.” (http://cr.yp.to/bib/imu-call.html). Again, one of the most important reasons for OA adoption is that it is recognised as an important tool to guarantee free access to scientific literature, considering that taxpayers pay for most of the research through government grants, and they must have the right to visualize to the results of what they have funded. Finally, it has also been suggested that scientists should benefit from the OA adoption because it could assure an increase in the impact of their research by potentially increasing the number of research papers citations (the so called “citational advantage” of OA) ^2^. This last point, in particular, has attracted the attention of researchers because the evaluation of research outputs impact, assessed as number of citations, is a consolidated practice as demonstrated by the popularity of citation-dependent bibliometric indexes (i.e. H index, IF, SJR, ect…) ^3,4^. It is often used in contexts related with the funding research policy, with the developing of carriers, as well as with the scientific collaboration policy ^5^. Consequently, it has a deep impact on scientists as well as on the public at large. However, the concept of OA citation advantage is still debated and there are conflicting hypotheses on this issue due to the complexity of the matter ^6,7^. For instance, there are several interfering factors that could affect the number of citations and are correlated to the OA citations without a causal factor such as the number of co-authors, Authors self-selection of higher quality articles for OA (Quality or Selection bias), the earlier dissemination via preprints/OA repositories (Early access bias) (http://www.istl.org/10-winter/article2.html). In that context, we performed an analysis (based on quantitative bibliometric data) aimed to study the dynamics related to the temporal and geographical spreading of OA publishing and the impact (evaluated in terms of number of documents, number of citations *per* document, SJR and H index) of medical scientific literature published under NOA or OA conditions. For this purpose, we accessed the data indexed in Scopus, which is a repository that includes Journals that fulfil basic quality requirements, thus avoiding to consider lower quality publications. We referred our analysis to medicine because the access, spreading, and impact of scientific knowledge in this field could have very important issues either ethical ^8,9^ or related to the data quality ^10^.

## Materials and Methods

### Data collection

As data source, we selected the Journals indexed on Scimago Journal and Country Rank (https://www.scimagojr.com/index.php) database. We selected this database because it assures an “*a priori”* quality control of indexed Journals. In fact, to be included in Scopus Journals, it is necessary to meet some eligibility criteria that assure a minimum of quality. In particular, they should consist of peer-reviewed content; they must be published on a regular basis and have an ISSN number registered with the International ISSN Centre. Furthermore, their content should be relevant and readable for an international audience, and the Journals should have a publication ethics and publication malpractice statement. Additionally, Journals should have a publication history of at least two years ^11^. If a Journal meets all these criteria, after an accurate peer-review process, it is indexed. Thus, a myriad of low quality Journals are excluded from our analysis. Indeed, mostly in recent years, we are seeing the proliferation of OA Journals with no regulation and scientific quality, often as part of the so called predatory publishing ^12,13^. Since in the present study, we were interested in assessing the impact of OA on the real scientific community, we analysed only the Journals present in Scopus (which is at least considered a necessary, but not completely sufficient requisite to be considered reliable). Clearly, this implies a limitation in our analysis, due to the fact that Scopus covers less than half of the biomedical Journals listed in Ulrich’s periodical classified as ‘‘Academic/Scholarly’’^14^.

We used SCImago because it is a publicly available portal, thus it guarantees free access to data, for proofing or further analyses. In addition, it includes the indicators developed from the information contained in the Scopus database (Elsevier B.V.) related to scientific Journals grouped according to subject area (27 major thematic areas), subject category (313 specific subject categories) or country. Citation data is drawn from over 34,100 titles from more than 5,000 international publishers and country performance metrics from 239 countries worldwide. Importantly, it assures a quality control of the Journals listed (see Discussion section).

We carried out our analysis considering the data referring to the Subject Area (SA): “Medicine”; Subject Category: “All subject categories”; years: 2001, 2006, 2011, 2016.

We obtained a first list of all the Journals (AJ), and then using the option “Only Open Access Journals”, we obtained the list of OA Journals. Finally, we computed the list of non OA (NOA) Journals by subtracting the OA Journals from the AJ list.

For each Journal and each year, we listed the following parameters:

- SCImago Journal Rank (SJR): used by SCImago as a measure of scientific influence of scientific Journals. It accounts for both the number of citations received by the Journal and the importance of the Journals such citations come from. It expresses the average number of weighted citations received in the selected year by the documents published in the selected journal in the three previous years, i.e. weighted citations received in year X to documents published in the journal in years X-1, X-2 and X-3.
- *H* index: a researcher with an *h* index has published *h* papers, each of which has been cited in other papers at least *h* times. It is a metric that attempts to measure the impact of a Journal on the scientific community;
- Cites per Document (2 years): defined as the average citations per document in a 2 years period. It is computed considering the number of citations received by a journal in the current year to the documents published in the two previous years, i.e. citations received in year X to documents published in years X-1 and X-2.
- Country: the Country where the Journal has been published, i.e. where the Publisher has its legal address. This could lead to some imprecision, since it is possible that the location of publishers office is not related to the
- Journal area of interest. Consequently, this issue could in part bias the discussion of this specific point.

For an extensive explanation of the parameters used, see https://www.scimagojr.com/help.php.

### Geolocalization

Data related to the selected papers were processed for geospatial analysis by Sci^2^ Tool (Sci^2^ Team). We generated a visualization of the choropleth map that shows the geographic distribution of the NOA and OA Journals distinguished by shades of colour for each Country, proportional to the number of Journals published.

### Data analysis

The data were checked for normalcy by using the D’Agostino and Pearson test, which computes the skewness and kurtosis of data under analysis and quantifies how far the distribution is from the Gaussian distribution. As a result, it computes a single P value from the addition of these discrepancies.

Then, we processed the data as normal or not-normal, following the needs (see figure legends for the specific test used in each analysis). All the calculations have been performed with Past3 or Microsoft Excel 2013. The bibliometric data analysis on Subject Categories was carried out when we found a number of records >5, to assure a sufficient quality of the statistical analysis. In particular, we compared the data referred to the NOA and OA Journals, in terms of number of Journal, published in 2001, 2006, 2011, and 2016, Country of origin, and bibliometric indexes (SJR, H index, number of citations/Document). In respect to this last analysis, we computed the best curve fitting comparing different models (e.g. linear, exponential, power law, logarithmic, polynomial models) and we assessed the best fitting based on the highest coefficient of determination R^2^.

## Results

### Temporal and geographical trends in NOA and OA publishing in medicine

As shown in Figure 1, it is evident that in comparison to the examined period, the number of NOA Journals increased less than 10%, reaching a plateau of about 5200-5300 Journals. The OA, on the contrary, are strongly increasing (four-time increase in the last 15 years) with a markedly increasing trend.

**Figure 1:**
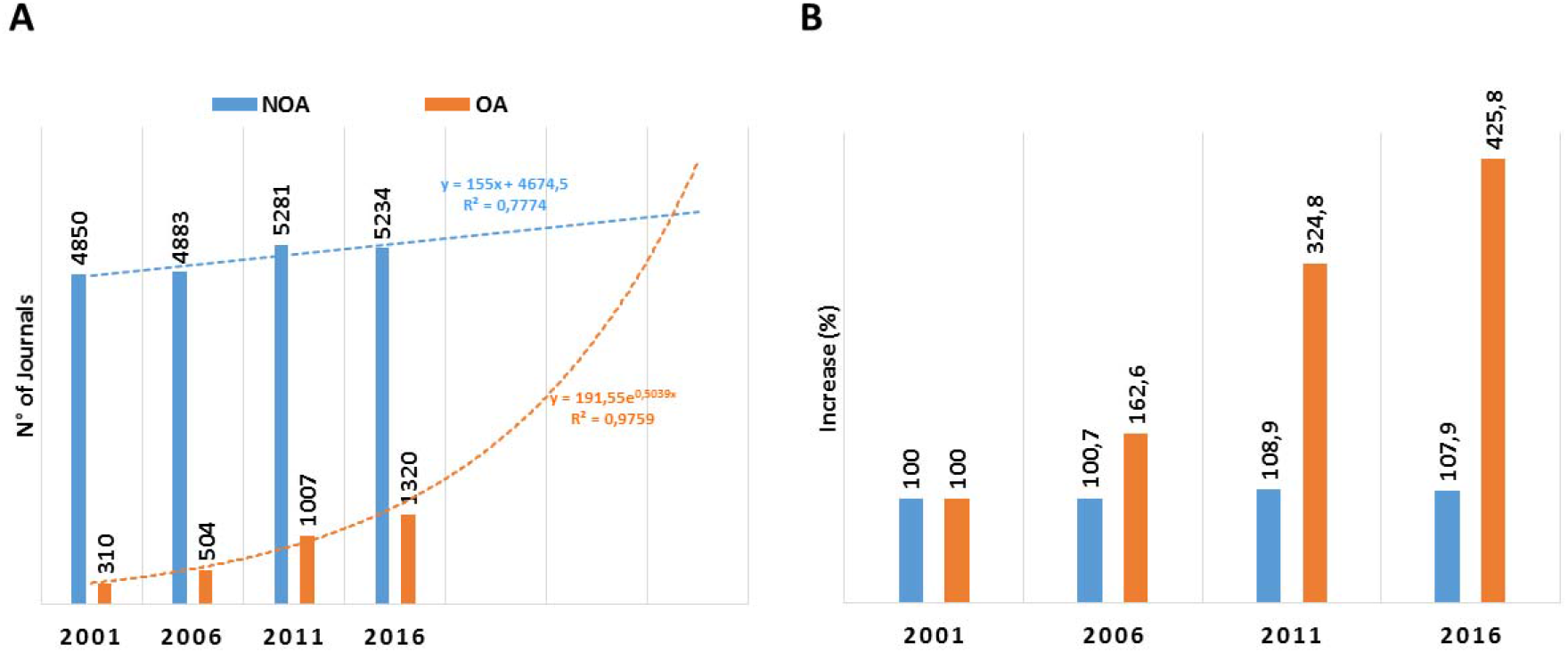
number of NOA and OA Journals in “Medicine” in 2001, 2006, 2011, and 2016 (Scopus). Data shown the number of Journals in each year as absolute number (Panel A) and as percentage (the value in 2001 = 100%) (Panel B). The light blue and orange dot lines represent the temporal trend in NOA and OA publishing of NOA and OA Journals, respectively.

We carried out the Geolocalization of NOA and OA Journals in accord with the period taken in examination. As a result, we computed the number of NOA and OA Journals active in 2001, 2006, 2011 and 2016. Data are listed in Supplementary Material 2 and graphically represented in Figures 2A-D.

**Figure 2:**
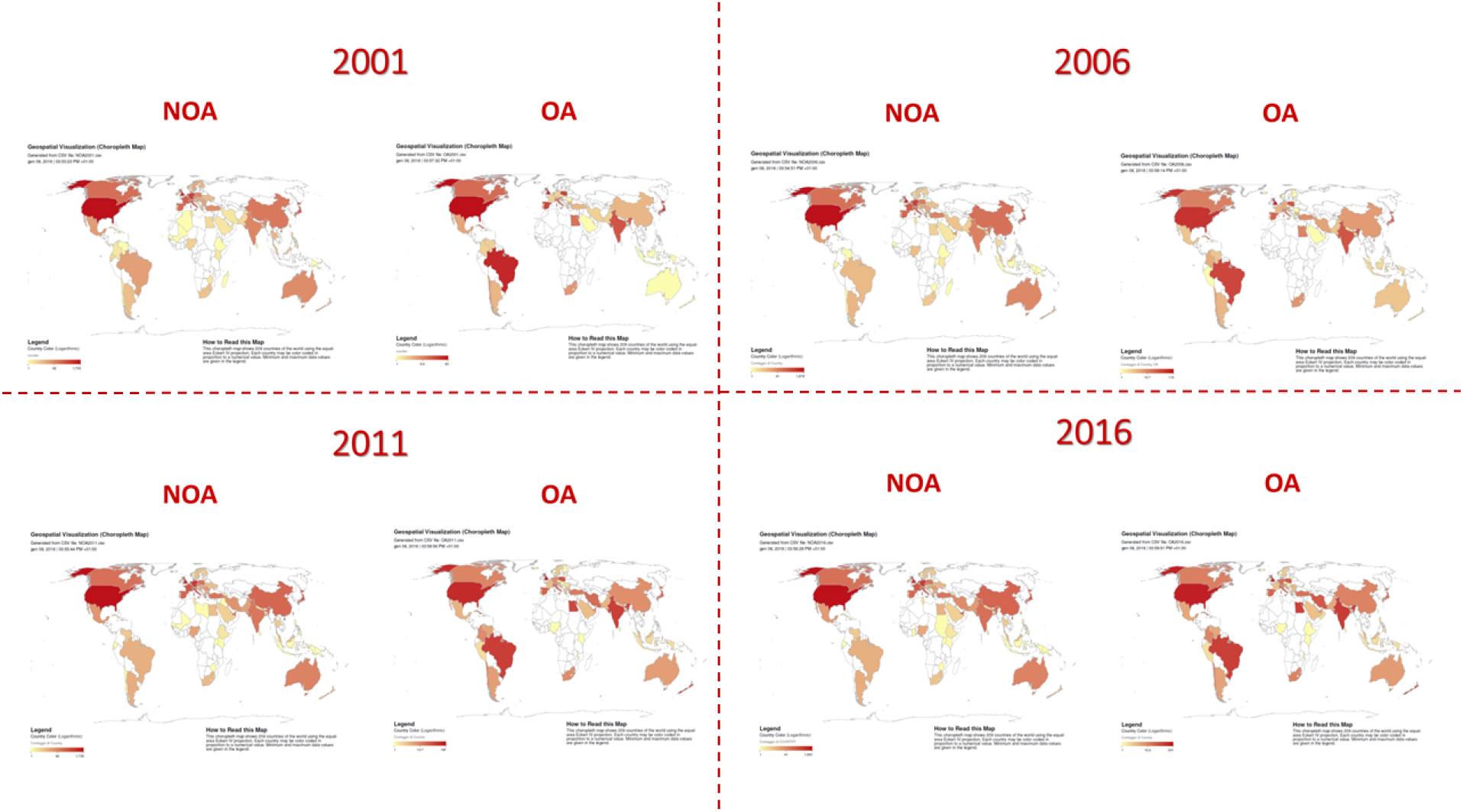
choropleth map of NOA and OA Publishers of Journals in “Medicine” in 2001, 2006, 2011, and 2016 (Scopus). The reported choropleth map shows the geographic distribution of the NOA and OA Journals differentiated by shades of colour for each Country on the basis of the number of published Journal.

### Could OA publishing assure a higher impact of research output in medical field?

As impact parameters, we evaluated SJR, H index and the number of Cites/Documents, referred to the years 2001, 2006, 2011 and 2016.

As it is evident in Figures 3, 4 and 5, all these parameters follow an exponential distribution, with a negative exponent ranging between 2 and 3.

**Figure 3:**
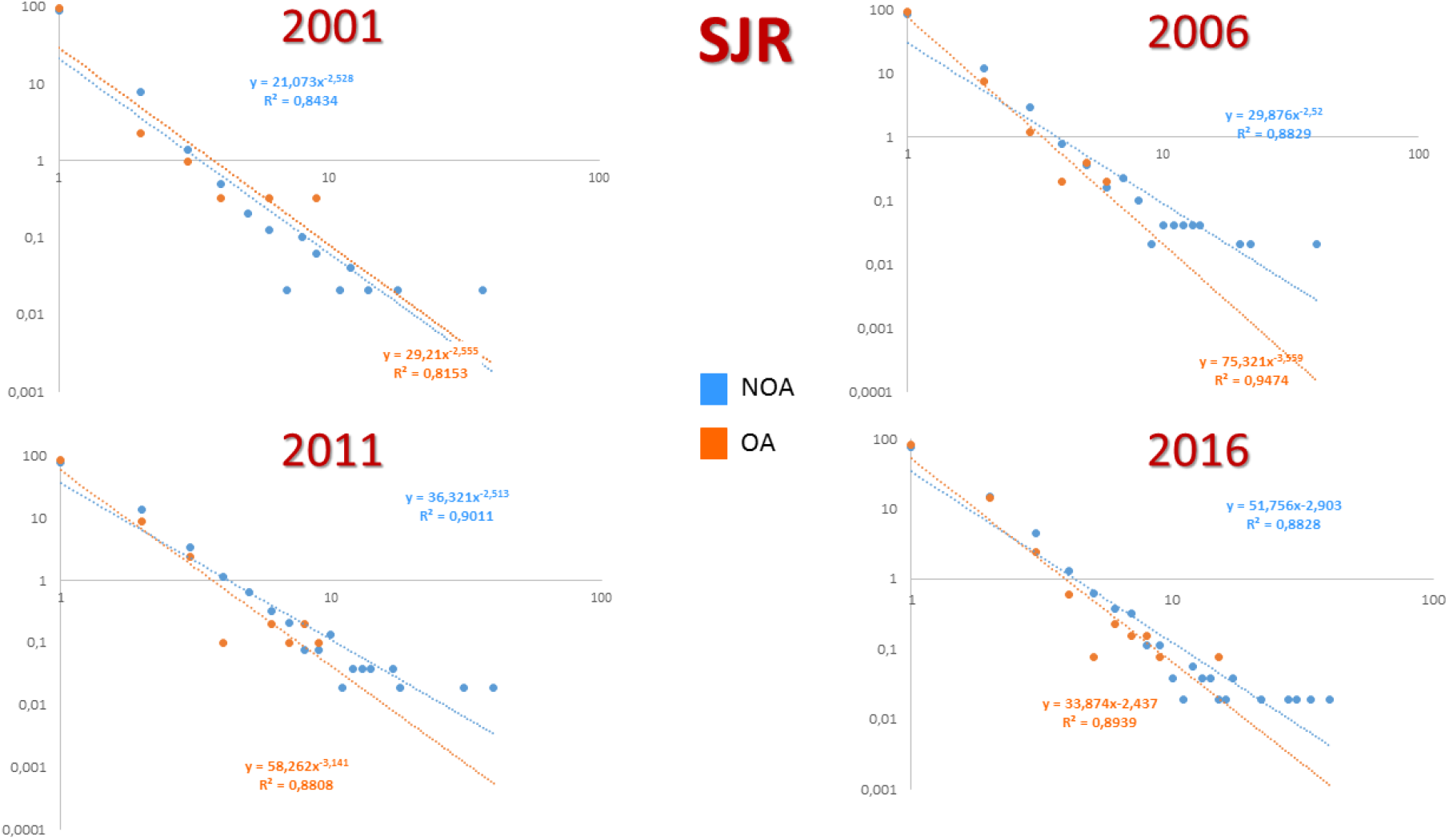
graphs showing the distribution of SJR in 2001, 2006, 2011, and 2016 in NOA and OA Journals.

**Figure 4:**
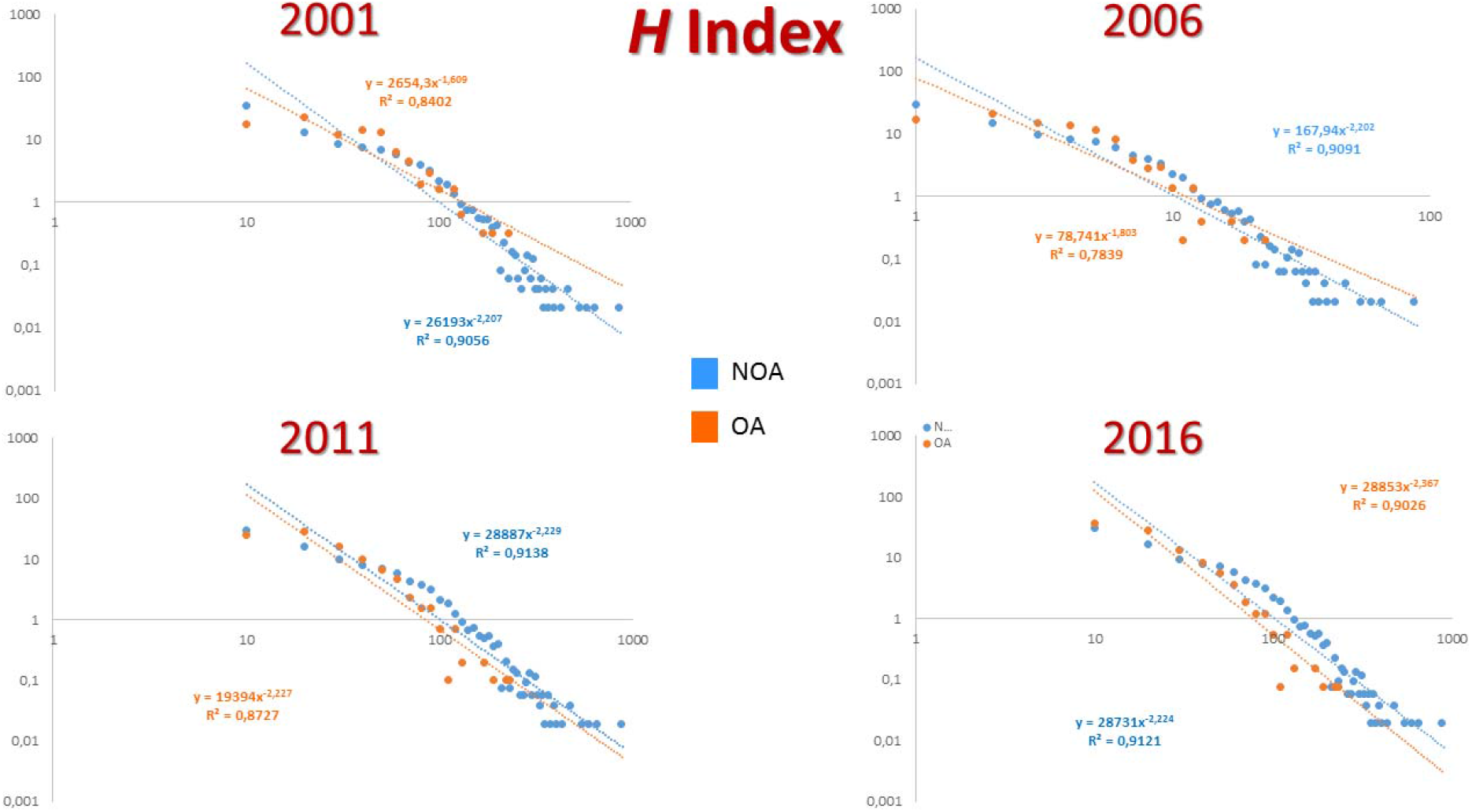
graphs showing the distribution of Cites/Document in 2001, 2006, 2011, and 2016 in NOA and OA Journals.

**Figure 5:**
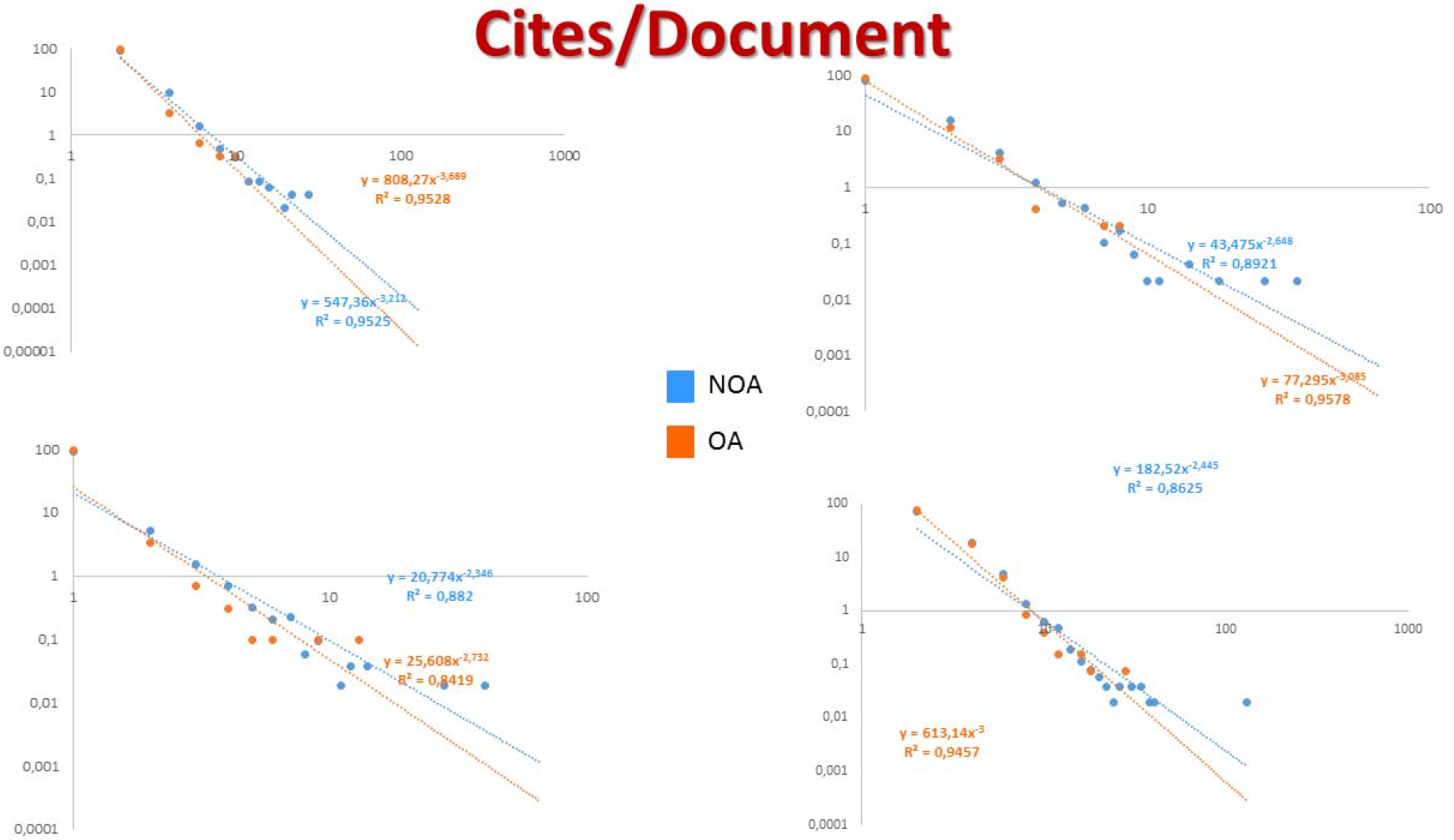
graphs showing the distribution of H index in 2001, 2006, 2011, and 2016 in NOA and OA Journals.

Interestingly, the exponents of SJR and Cites/Document are always lower in OA when compared with NOA, while the time trend seems to be stable. On the contrary, H index was higher in OA in 2001, then it gradually decreased until 2016, when it was higher than that of NOA Journals, as displayed in Figure 6.

**Figure 6:**
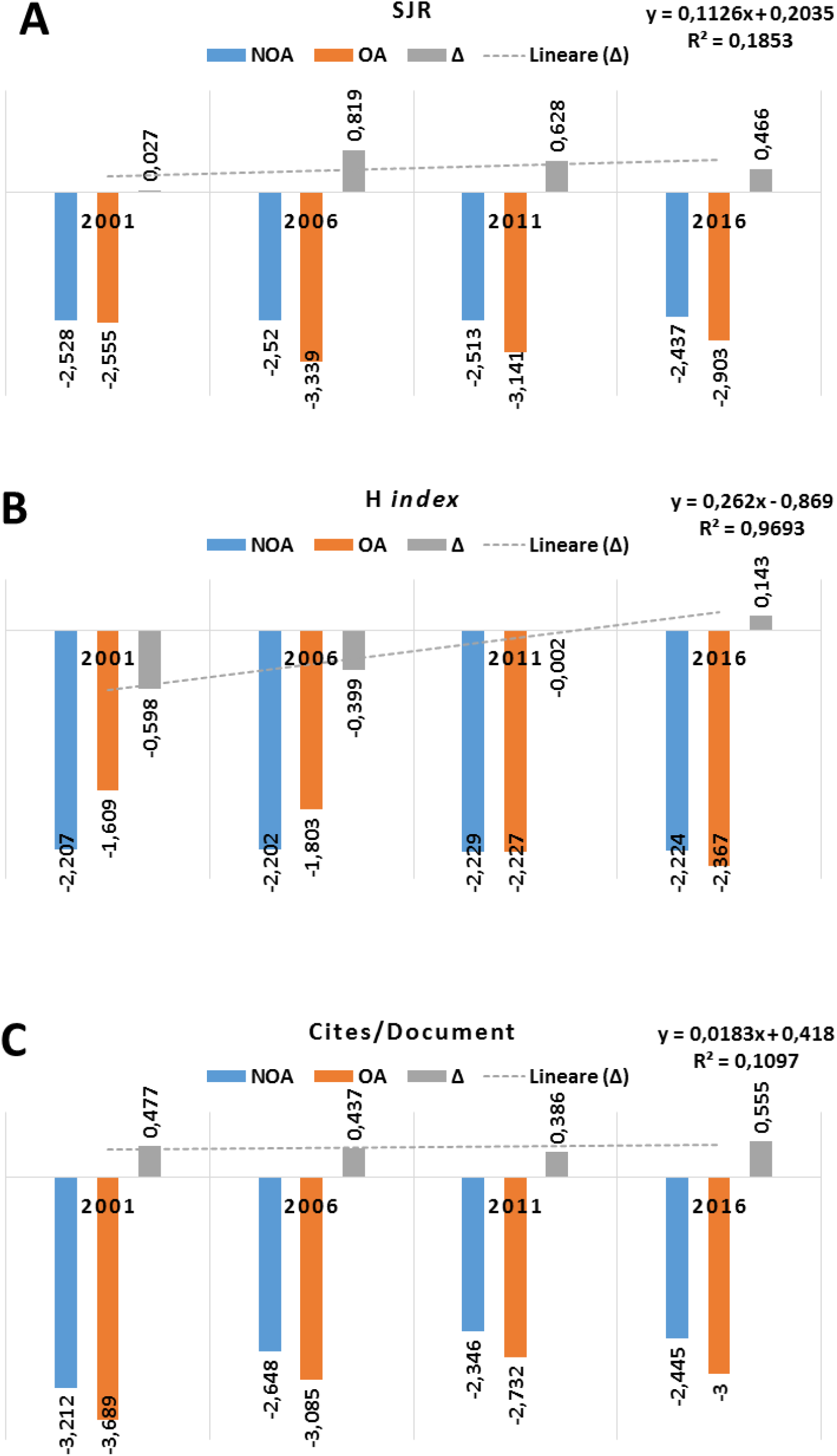
graphs showing the exonents of bibliometric parametrs (SJR, H index, and Cites/Document) in 2001, 2006, 2011, and 2016 in NOA and OA Journals.

The differences in term of bibliometric indexes are summarized in Table 1.

**Table 1:** list of *p* values obtained by comparing NOA and OA Journals (data compared with Mann–Whitney *U* test)

| Year | SJR | H Index | Cites/Document (2 years) |
| --- | --- | --- | --- |
| 2001 | <0,0001 | 0,0037 | 0,0207 |
| 2006 | 0,0333 | 0,2052 | 0,9653 |
| 2011 | 0,0037 | <0,0001 | 0,8190 |
| 2016 | 0,0733 | <0,0001 | <0,0001 |

Were $p < 0.05$ NOA are always higher than OA.

As summarized in Supporting Material 1, we analysed all bibliometric parameters studied in all Subject Categories, with a minimum number of 5 Journals. On a total of 465 parameters, we found that in about 82% of cases (381/465) there aren’t differences. While in 18% (84/465) we found statistically significant differences (p<0.05). In all cases, the impact parameters referred to NOA were higher than those of OA Journals were.

## Discussion

Here, for the first time to our knowledge, we carried out an analysis aimed to take an updated picture of OA adoption in publishing in Journals in the medical field using Scopus as data source (https://www.scopus.com/). In particular, we evaluated the number of NOA and OA Journals in medicine over the time (2001-2016). Evidently, the number of NOA Journals increased less than 10% in the last 15 years and seems to have reached a plateau. On the contrary, the OA Journals in 2001 were 310 (vs. 4580 NOA Journals) while in 2016 they were 1320, with an increase of over 400%, furthermore the trend seems to be continuously increasing with an exponential trend (191.6 ; R^2^ = 0.976). Although these data are only indicative, it is possible to affirm that the OA model seems to be a very successful, so that if the trend does not change in the future it could be possible that in less than 15 years, the number of OA Journals will overcome that of NOA Journals. See Figure 1. Consequently, we analysed the data related to the most productive Countries, responsible for the increasing number of OA products. In this regard, it is very interesting to note that the Countries mostly active in the increasing of number OA Journals are Brazil, India, South Africa, New Zealand, Poland, Serbia, and South Korea, while the Countries with a leader position and a well-defined scientific tradition (USA, UK) did not display a comparable trend. From this datum, we can infer that OA could open new markets to the publishing system. Interestingly, three of them are BRICS (Brazil, Russia, India, China and South Africa) members. They are large and fast-growing national economies, characterized by specific demographic and geographic situations that play a specific and leading role with a significant influence on regional and global affairs. In 2015, they represented 3.6 billion people, 22% of the world economy (gross world product) and nearly 60% of its growth ^18^. Brazil today is the leading Country in Latin America, with more than 1,700 journals alone on the Open Journal System. In 2015, 36% of the Brazilian research output was open access (compared to 12-14% for France, Germany, US or UK) ^19,20^. The most important OA platform in Brazil is the Scientific Electronic Library On-line, SciELO (http://www.scielo.br/). It is “an electronic library covering a selected collection of Brazilian scientific journals” (http://www.scielo.br/), and it is integral part of a project developed by - Fundação de Amparo à Pesquisa do Estado de São Paulo (FAPESP), in partnership with the Latin American and Caribbean Center on Health Sciences Information (BIREME), supported by - Conselho Nacional de Desenvolvimento Científico e Tecnológico (CNPq). The main aim of the project is “the development of a common methodology for the preparation, storage, dissemination and evaluation of scientific literature in electronic format” (http://www.scielo.br/). This has had the great advantage to make available, without costs, the research results to the researchers and lay people, mainly in Latin America. But, unfortunately, as criticized, without considering the quality of Journals indexed ^18^. To date (February 2018) it accounts more than 20,000 documents (82.6% research articles), mainly of Brazilian Authors (70.9%), of which more than half are in English (56.8%), with particular regard to Health Sciences (38.7%), Human Sciences (17.8%), and Agricultural Sciences (14.1%)(all the information related SCIelo is listed in Supporting Material 2).

In India, the OA policy is supported by important research organisations, laboratories and universities (10). Consequently, more than half of Indian Journals are characterized by OA publishing and most of them do not charge author fees ^21^. This, in one hand increased the availability and the visibility of Indian research product; on the other hand, it makes the way to the spread of predatory publishing. As reported by Beall (https://beallslist.weebly.com/) several Journals declare to be headquartered in the United States, United Kingdom, Canada or Australia, but are really located in Pakistan, Nigeria or, mainly, India.

South Africa has the second largest economy in Africa, after Nigeria, and it is Country more industrialized in Africa. It accounts for 35^%^ of Africa gross domestic product and is ranked as an upper-middle-income economy by the World Bank, and is the only African member of the G-20 group. Because of its technological infrastructures it has quickly developed online journals, repositories, and a variety of other tools and platforms based on OA, South Africa is the leading African Country in terms of OA policies ^22^. To date, the international Directory of Open Access Journals records 79 open access journals produced in South Africa (http://roar.eprints.org/). In South Korea the OA adoption is growing very rapidly in recent years and its improvement is part of the great effort in sustaining education and high technology research ^23^.

New Zealand Institutions are quickly moving towards adoption OA policies and mandates. For instance, contracts for the Marsden Fund, New Zealand’s fundamental research fund, include a clause mandating that Researchers must share their data, meta-data and samples within 12 months of completion of the project (https://aoasg.org.au/new-zealand-open-access-journals/).

The specific situation of OA publishing of Poland and Serbia, the only two European Countries examined, are exhaustively described in two monographs to which we refer the reader ^24,25^.

Then, we assessed three different parameters (SJR, H index and number of cites/document) that are widely used to evaluate the impact of Journals, related to citations. In particular, SJR assigns different values to citations depending on the importance of the journals and where they come from. Consequently, citations coming from important Journals, will be considered (weighted) more than those from less important Journals. The SJR calculation is similar to the Eigenfactor score in the Web of Science database ^15^. H index, suggested in 2005 by Jorge E. Hirsch, which represents the measure of impact and productivity of an Author, a group of Authors or a Journal, as well as the number of cites/document, related to the unweighted number of citations.

We are conscious that the use of citation-related indexes could be criticized for several valid reasons and that the relationship between the number of citations of a scientific paper and the importance of its content is not always obvious, and the values of the parameters used, are strongly dependent on the database accessed and on the specific discipline. For these reasons, we limited their use as indicators of Journal impact rather than Journal quality ^16,17^.

The comparison of bibliometric parameters between NOA and OA Journals demonstrates that they are often statistically different trough the examined years, with NOA Journals showing higher values.

More interesting, we compared them in the different subject categories (SC) of medical areas. Here too, we found several important differences, with the parameters of NOA Journals higher than those of OA ones in about 20% of the cases (85 of 464).

Another interesting thing is the study of frequency distribution of these indexes. Indeed, in agreement with the Bedford’s law, they are distributed following a power law, with a negative exponent (usually near 3). As it is evident from Figure 6, for SJR and Cites/Document parameters the exponent of NOA Journals is virtually always higher (lower in absolute value), with a stable trend over the time. It means that that the distribution of values is flatter. In other words, a higher number of Journals (in proportion) have lower values and a lower number of Journals has higher values. These results seem to be in contradiction with the previous results from Eysenbach that found a “Citation Advantage of Open Access Articles” ^2^. Actually, he compared articles published in the same Journal (PNAS), which adopted a hybrid publishing policy (it means that part of articles were published with NOA and part with OA). Adopting a logistic regression model, controlling for potential confounders, he found that OA articles compared to NOA articles remained twice as likely to be cited in the first 4–10 months after publication (odds ratio= 2.1 [1.5–2.9]), with the odds ratio increasing to 2.9 (1.5–5.5) 10–16 months after publication. Then, it was concluded that “Articles published as an immediate OA article on the journal site have higher impact than self-archived or otherwise openly accessible OA articles.” ^2^ In other words, it was suggested that the adoption of OA could assure a higher impact on the scientific community, when compared to NOA.

In our opinion, the apparent difference with our results is due to the different rationale of the investigation. Eysenbach compared the citations of articles published in the same Journal, PNAS, which is a very peculiar Journal: it is a general science journal, with an H-index = 648, and an SJR = 6.321 (2016) ^2^. While this study is focused on PNAS, which is a Journal with very peculiar characteristics, we compared a very large number of Journals referred to whole the medical area, consequently his conclusion are not directly comparable with those from our data analysis.

Anyway, the question remains open, as proven by the conclusion of another very interesting work on PNAS citations of NOA and OA papers from Patrick Gaulé and Nicolas Maystre ^12^. In fact, they concluded that “at least part of the large number of citations received by open access papers is due to a self-selection effect rather than a spreading (or causal) effect”. In means, that Authors preferentially published as OA the articles they consider of higher quality and the reason of the citation number could actually be the quality itself. As a proof, they found that OA articles receive a higher number of cumulative citations also in articles published in Nature, Science, or Cell, that are authored by scientists performing cutting-edge science and that can hardly be expected to lack access to the NOA scientific literature.

Further, the Open Access Citation Advantage Service (OACA), which is a non-profit, member organisation comprised of a diverse body of academic institutions, library consortia, funding bodies, research institutes and some publishers published on its website (https://sparceurope.org/) a list of 70 articles until 2015, of which 66% (46) declare a citation advantage for OA papers, 24% (17) found no advantages and 10% (7) were inconclusive. Unfortunately, the inclusion criteria of the articles and the quality check methods were not discussed, and then this data cannot be considered actually reliable.

Ultimately, the relevance of OA in sharing research results among scientists and/or lay people is challenged by different ways of sharing. In particular, widely accessed repositories are often used, as for instance ResearchGate (https://www.researchgate.net/), as well as other platforms (SciHub, OpenAccessButton, Unpaywall); the assessment of their impact is out of the scope of this paper, see for reference the comment: “Who’s downloading pirated papers? Everyone.” by John Bohannon (http://www.sciencemag.org/news/2016/04/whos-downloading-pirated-papers-everyone).

## Conclusions

Despite this study has limitation due to the fact that Scopus only distinguishes between OA and non-OA journals, and not OA and non-OA articles thus the analysis inevitably is unable to account for articles published OA in hybrid journals.

Anyway, we think that our analysis could contribute to the knowledge of the OA model spreading in the medical field, offering a new perspective since it takes into account the Journals listed in Scopus referable to Medicine published in a large window of time (2001-2016) that coincides with the onset and spreading of OA literature. In conclusion, we could affirm that:

- We are seeing in medicine a huge increase in the number of Journals that use OA;
- This trend is mainly due to the increase in publishing Journals in Countries with emergent economies (Brazil, India, South Africa, New Zealand, Poland, Serbia, and South Korea), rather than in developed occidental economies;
- The use of OA for Journals in the medical field, in the last 15 years did not guarantee a vantage in term of impact, but rather they seem often to be characterized by lower impact parameters.

In 1999, The World Conference on Science held under the auspices of UNESCO and ICSU stated that “Equal access to science is not only a social and ethical requirement for human development, but also essential for realizing the full potential of scientific communities worldwide and for orienting scientific progress towards meeting the needs of humankind” ^26^. We could conclude, after almost 20 years, that researchers of the emerging countries significantly adopt OA. Even if, nowadays, it does not seem able to ensure a higher impact in publication in the medical field, it represents a new and rapidly expanding way to share medical knowledge.

## Authors Contribution

NB: Conception and design of the work, analysis, and interpretation of data, writing of manuscript; RC and AO: acquisition and analysis of data, contributing in interpretation of data; BB and LV: revising the article critically for important intellectual content and for interpretation of data; All: final approval of the version to be published and agreement to be accountable for all aspects of the work.

